# Spatial-temporal expression analysis of lineage-restricted shell matrix proteins reveals shell field regionalization and distinct cell populations in the slipper snail *Crepidula atrasolea*

**DOI:** 10.1101/2023.03.18.532128

**Authors:** Rebecca N. Lopez-Anido, Grant O. Batzel, Gabriela Ramirez, Jessica A. Goodheart, Yiqun Wang, Stephanie Neal, Deirdre C. Lyons

## Abstract

Molluscs are one of the most morphologically diverse clades of metazoans, exhibiting an immense diversification of calcium carbonate structures, such as the shell. Biomineralization of the calcified shell is dependent on shell matrix proteins (SMPs). While SMP diversity is hypothesized to drive molluscan shell diversity, we are just starting to unravel SMP evolutionary history and biology. Here we leveraged two complementary model mollusc systems, *Crepidula fornicata* and *Crepidula atrasolea*, to determine the lineage-specificity of 185 *Crepidula* SMPs. We found that 95% of the adult *C. fornicata* shell proteome belongs to conserved metazoan and molluscan orthogroups, with molluscan-restricted orthogroups containing half of all SMPs in the shell proteome. The low number of *C. fornicata*-restricted SMPs contradicts the generally-held notion that an animal’s biomineralization toolkit is dominated by mostly novel genes. Next, we selected a subset of lineage-restricted SMPs for spatial-temporal analysis using *in situ* hybridization chain reaction (HCR) during larval stages in *C. atrasolea*. We found that 12 out of 18 SMPs analyzed are expressed in the shell field. Notably, these genes are present in 5 expression patterns, which define at least three distinct cell populations within the shell field. These results represent the most comprehensive analysis of gastropod SMP evolutionary age and shell field expression patterns to date. Collectively, these data lay the foundation for future work to interrogate the molecular mechanisms and cell fate decisions underlying molluscan mantle specification and diversification.

## Introduction

Biomineralization is a significant innovation in the evolution of divergent body forms (Gilbert et al. 2022), and is widespread throughout the Metazoa (Fig. 1A). Within the phylum Mollusca there exists a rich diversity of biomineralized shells with different shapes, sizes, colors, microstructures, and mineral content (Fig. 1A) (Chateigner et al. 2000; McDougall et al. 2013; Kocot et al. 2016; Williams 2017). Shells are built through the process of biomineralization, during which specialized epithelial cells in the mantle tissue secrete organic materials, including lipids, glycoproteins, and shell matrix proteins (SMPs), into an enclosed, extrapallial space. Secreted SMPs self-assemble to create an extracellular matrix that facilitates the precipitation of calcium carbonate (Falini et al. 1996; Thompson et al. 2000; Xie et al. 2016). In the field of molluscan biomineralization, SMP diversity is credited as the driving force behind novel shell morphologies (Kocot et al. 2016).

**Figure 1.**
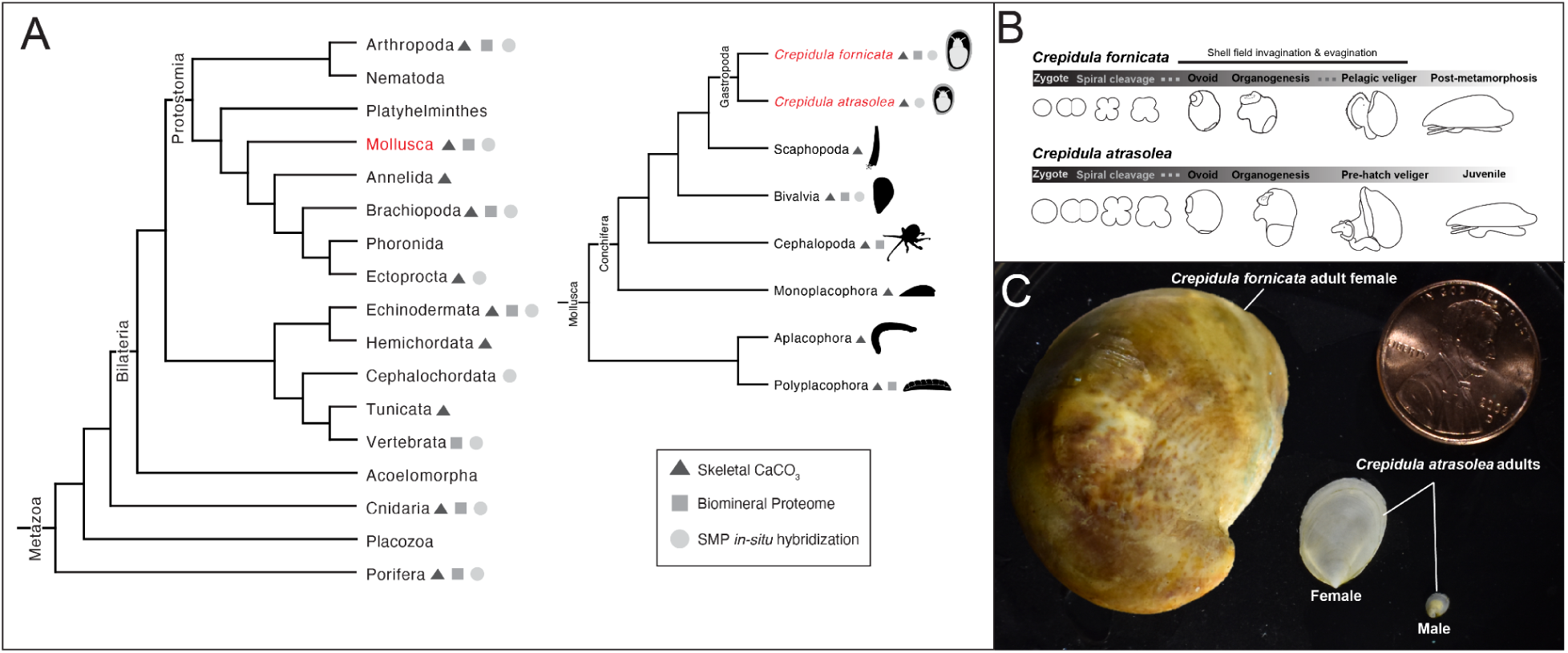
*Crepidula fornicata* and *Crepidula atrasolea* are complementary model systems for biomineralization. Phylogeny of calcium carbonate biomineralization throughout the Metazoa (left) (Gilbert et al. 2022) and the Mollusca (right) (Kocot et al. 2020). Triangles indicate taxa that produce calcium carbonate skeletons; squares indicate taxa that have a published biomineral proteome; circles indicate taxa that have conducted *in-situ* hybridization for shell matrix proteins (SMPs) (A). Comparison of indirect larval development in *C. fornicata* vs direct development in *C. atrasolea* (C). Adult shells of the two species (D).

Our current understanding of SMP diversity is largely based on proteomic investigations that have identified SMPs occluded in the shells of different molluscan species (Marin 2020). Sequence similarity searches of these SMPs have been used to help annotate and assess the evolutionary origins of SMPs, and through this work, many species-restricted SMPs have been identified (Song et al. 2019). For example, studies of bivalve and gastropod shell proteomes routinely report high percentages of species-restricted SMPs (also referred to as “novel” or “orphan” genes), and a smaller percentage of conserved SMPs (Clark 2020). These homology-based classifications are traditionally determined by BLAST searches of SMPs against large sequence databases like GenBank and NCBI; if SMPs have no best-reciprocal BLAST hit when searching these databases they are classified as species-restricted (as they are found only in the species being searched), whereas those with reciprocal BLAST hits are placed at the taxonomic level they are restricted to (Weisman et al. 2020). SMPs shared by metazoans are often found to be deeply conserved genes involved in fundamental processes like calcium regulation (e.g. calmodulin) or bicarbonate production (e.g. carbonic anhydrase) that have been co-opted for biomineralization functions (Means and Dedman 1980; Livingston et al. 2006; Le Roy et al. 2015). In contrast, lineage-restricted SMPs are viewed as fast-evolving and recently-diverging genes, with repetitive, low complexity regions in their gene sequences (McDougall et al. 2013). Thus, the evolutionary age of an SMP may dictate its functional role in shell morphological characteristics, with conserved SMPs providing the basic physiological properties of a shell (precipitation of minerals), while recently evolved lineage-restricted genes contribute to novel shell phenotypes. Examples of lineage-restricted SMPs potentially driving novel shell phenotypes include SMPs influencing pigmentation in gastropods (Mann and Jackson 2014) or regulating shell nacre production in bivalves (Suzuki et al. 2009). In addition to SMP lineage-specificity, spatial expression of SMPs within the biomineralizing mantle might be a second factor that determines novel shell phenotypes. For example, in adult bivalves, SMPs involved in nacreous-layer formation are found in the dorsal region of the mantle epithelium, whereas SMPs involved in prismatic-layer formation are localized to the ventral portion of the mantle epithelium (Takeuchi and Endo 2006).

However, there is still much more to be gleaned by placing molluscan modularity within an evolutionary context, to answer the extent to which lineage-restricted SMPs and their cell type specific expression contribute to either novel or conserved shell morphological characteristics. These questions are best addressed during larval shell development, a stage when lineage-restricted SMPs are first expressed, and when mantle cell differentiation and morphogenesis are occurring.

In most gastropods, shell development begins after gastrulation when the dorsal epithelium thickens and becomes known as the shell field (Fig. 1B). During larval development, the dorsal shell field epithelium invaginates to produce a transient gland structure, known as the shell gland (Kniprath 1981; Eyster and Morse 1984), where organic shell secretions are first detected (Kniprath 1981; Hohagen and Jackson 2013). The shell gland disappears when the shell field subsequently evaginates to form the larval mantle epithelium. The larval mantle epithelium continues to secrete shell material, including SMPs, throughout larval development and into adulthood. In the literature, “shell field” and “larval mantle epithelium” both refer to the larval tissues responsible for shell formation. Larval SMP expression patterns have been examined in molluscan species (Werner et al. 2013; Gaume et al. 2014; Herlitze et al. 2018; Batzel et al. 2022), but with minimal cellular resolution due to the limitations of traditional, colorimetric *in-situ* hybridization. To detect additional SMP expression patterns and SMP-expressing cell populations, a finer cellular and subcellular resolution of signal is required.

In this study we leveraged two species in the *Crepidula* genus of slipper snails, *Crepidula fornicata* and *Crepidula atrasolea* (Fig. 1A: Fig. 1C), to present the first systematic study of SMP lineage-specificity and spatial-temporal expression during shell development. First, we establish *C. fornicata* and *C. atrasolea* as complementary models for studying molluscan biomineralization. We used orthology inference to determine the evolutionary age and level of lineage-restriction of 185 SMPs from the adult shell of *C. fornicata* (Batzel et al. 2022). Next, we examined the larval expression patterns of 18 conserved SMPs in *C. atrasolea* using hybridization chain reaction (HCR) and confocal imaging. Lastly, we characterize potential regions of larval biomineralization, and identify shell field cell populations. Together, we consider the relationship between SMP evolutionary age and shell field regionalization to provide new perspectives on molluscan shell diversification.

## Methods

### Animal culture

*Crepidula atrasolea* (Collin 2000) adults were maintained in the Lyons lab at the Scripps Institution of Oceanography, San Diego, California, USA. The animals were kept at 27°C, in 100 mm x 20 mm plastic petri dishes (Corning # 430167) with daily filtered natural seawater changes, and fed daily with 0.002% Phyto-Feast (Reed Mariculture Inc., Campbell, CA). Clutches of eggs inside their capsules were collected from the adults using forceps (as previously described in Henry et al. 2017) and placed in 35 mm x 10 mm plastic petri dishes (Falcon # 351007) with bottle top-filtered (0.2 μm pore size; Thermo Scientific # 291-4520) natural sea water with streptomycin (2ug/mL; Sigma # S6501-100G) and penicillin (0.6 ug/mL; Sigma # 13752-5G-F). The collected embryos were then raised at room temperature (∼20°C) inside their capsules until reaching the appropriate stages needed for fixation. *C. atrasolea* eggs are fertilized internally, and thus the timing of fertilization is based on hours or days since the embryo was uncleaved. The development of embryos within a single clutch is asynchronous, resulting in uncleaved, 2-cell, and 4-cell embryos all in the same clutch. In addition to days post fertilization (dpf), we examined the morphology of the developing mouth, velar lobes, foot rudiment, and shell field to stage older embryos. When embryos reached a desired stage, they were decapsulated under a dissecting scope using forceps in a gelatin coated 35 mm x 10 mm plastic petri dish prior to fixation.

### Embryo fixation

For embryos with ciliated velar lobes (starting at the organogenesis stage; Fig 1C), individuals were relaxed in 7.5% magnesium chloride (dissolved in filtered natural seawater; Sigma-Aldrich #M7304) at benchtop for 30 minutes prior to fixation. Decapsulated embryos were fixed at ∼20°C with 4% formaldehyde (Thermo Scientific # 28908) in double filtered sea water for 60 minutes on a shaking plate followed by three 1 minute 1X PBS Tween (1X Phosphate Buffered Saline: Gibco #70011-044, 0.5% Tween 20: Sigma-Aldrich P1379-100 mL) washes followed by three 10 minute 1X PBS Tween washes. Animals were then dehydrated in methanol with three 5 minute 100% methanol washes followed by three 1 minute 100% methanol washes. Fixed individuals were stored at −80°C in 100% methanol.

### Wheat Germ Agglutinin

Selected samples fixed in methanol (detailed above) were rehydrated into 5X SSCT solution (5X sodium chloride sodium citrate: Invitrogen # 15557044, 0.1% Tween 20) following 5 minute 75% methanol 25% 5X SSCT, 50% methanol 50% 5X SSCT, 25% methanol 75% 5X SSCT, and two 100% 5X SSCT washes. Embryos were then stained with 0.001 mg/ml wheat germ agglutinin (WGA, Alexa Fluor™ 594 Conjugate; Invitrogen W11262) overnight in the dark at 4°C, and rinsed after with the 5X SSCT solution three times for 5 minutes at 20°C. Samples were stored in the dark at 4°C, and imaged within a week. Imaging methods are detailed below.

### Sample collection and RNA extraction

#### Collection of embryonic stages

Three embryonic samples (Samples 1-3) spanning zygote through early juvenile development were collected for RNA extraction. The three embryonic staged samples each contain two pools of embryos that were kept separate for RNA extraction purposes, but later equal amounts of RNA were combined between pools to form Samples 1-3 for library preparation (see “RNA extraction” subsection for more details). Sample 1 contains one pool (n = 279) of uncleaved zygotes (0-8 hpf), 25-cell (24 hpf), and compaction (∼36 hpf) staged embryos; and a second pool (n = 186) of rectangular stage embryos (5 dpf). Sample 2 includes one pool (n = 180) of early organogenesis stage embryos (7 dpf) and a second pool (n = 168) of late organogenesis stage embryos (8-11 dpf). Sample 3 includes one pool (n = 133) of pre-hatch juveniles (14 dpf) and a second pool (n = 55) of post-hatched juveniles (14-15 dpf). Pools of embryos were kept separate and transferred to 1.5 mL tubes for RNA extraction.

#### Collection of adults and and their organs

Three samples (Sample 4-6) of *C. atrasolea* adults or their organs were collected for RNA extraction. Sample 4 contains one adult male specimen of *C. atrasolea*, while Samples 5-6 contain organs dissected from the same adult female. Whereas Sample 5 contains only the mantle, Sample 6 contains RNA from Sample 5 as well as the remaining organs (head, foot, gill, and visceral mass). Each dissected organ was kept separate for RNA extraction, and later recombined in equal parts to form Sample 6 prior to library preparation (see “RNA extraction” subsection for more details).

#### Dissection of organs

For organ dissection, a single adult female was placed into a 100 x 20 mm petri dish (Corning #430167) and relaxed in 7.5% magnesium chloride (dissolved in filtered natural seawater; Sigma-Aldrich #M7304) at room temperature for 30 minutes prior to undergoing dissection. After relaxation, fine forceps (Dumont #6, Fine Science Tools) were used to gently depress the foot from the shell deck followed by gentle pulling on the foot to remove the remaining organs found within the mantle cavity of the shell. Surgical scissors were used to remove the head organ at the intersection between the neck and the anterior margin of the foot. The anterior dorsal mantle (extending to around one third the length of the mantle cavity aperture) and the mantle border surrounding the foot, were dissected using both forceps and surgical scissors. The foot and visceral mass were separated from one another at their intersection between shell deck and inner mantle cavity. All four organs (mantle, foot, head, and visceral mass) were each placed into 1.5 ml tubes for RNA extraction.

#### RNA extraction

Total RNA was extracted using TRIzol (ThermoFisher), according to the manufacturer instructions (Invitrogen Doc. Part No. 15596026.PPS). Briefly, TRIzol was added to the collection tubes, and tubes were immediately frozen by placing them into a dry ice ethanol bath (70% ethanol, dry ice). Once frozen, mortar and pestle were used to completely homogenize tissues followed by the addition of chloroform to achieve separation of RNA in the aqueous layer. Precipitation of RNA from the aqueous layer was performed using 100% isopropanol, and after centrifugation, the RNA pellet was cleaned of impurities using 70% ethanol followed by resuspension in nuclease free water. Quantity and quality of RNA was conducted using a NanoDrop spectrophotometer. In total, 1 µg of RNA was collected for each of the six samples (Samples 1-6). For embryonic samples (Samples 1-3), 0.5 µg of RNA was combined from each of their respective two pools of embryos. For Sample 4, 1 ug of RNA was taken from the adult male. For Sample 5, 1 µg of RNA was taken from the mantle tissue. Lastly, for Sample 6, 250 µg of RNA each was taken from the head, foot, gill, visceral mass, and mantle.

### Iso-Seq RNA Sequencing

#### Library Preparation and Sequencing

Six RNA samples (described in the section above) were sent to the Roy J. Carver Biotechnology Center at the University of Illinois at Urbana-Champaign where the library preparation and sequencing were performed. 300 ng of total RNA were converted to cDNA with the Iso-Seq Express Oligo Kit. cDNAs were barcoded with the Barcoded Overhang Adaptor Kit 8A, and converted into a library with the SMRTBell Express Template Prep kit 2.0 (Pacific Biosciences). The libraries were pooled in equal concentration and the pool was sequenced on 1 SMRTcell 8M on a PacBio Sequel IIe using the CCS sequencing mode and a 30hs movie time.

#### Preprocessing and Clustering

The initial processing and clustering of the IsoSeq data was performed by Roy J. Carver Biotechnology Center at the University of Illinois at Urbana-Champaign using SMRTLink V10.1.0. Briefly, Circular Consensus Sequencing (CCS) analysis was first done using the “ccs” command with parameters “--min-passes 3 --min-rq 0.999”. Sample demultiplexing was then performed using the “lima” command with parameters “--ccs --same --split-bam-named”. This resulted in six bam files, each storing the reads from one of the six samples sequenced. Next, SMRTLink linux-toolkit V10.0.0 was used to 1. remove primers (lima --isoseq … primers.fasta …), 2. classify strandness and trim polyA (isoseq3 refine --require-polya), and 3. de novo cluster the transcripts (isoseq3 cluster --use-qvs) for each sample. The final output is a fasta file containing full length consensus isoforms for each sample. We additionally merged the bam files for all samples and repeated the clustering step using “isoseq3 cluster” command to generate a consensus isoform fasta file across all samples. This fasta file was used in the subsequent hybrid transcriptome construction.

### Hybrid transcriptome assembly and annotation

#### Obtaining short and long read transcriptome data

We used both short read transcriptomes generated by Illumina sequencing and long read transcriptomes generated by Iso-seq to construct a high coverage transcriptome. The long read transcriptomes were obtained as described in the previous section. The short read transcriptomes were previously published by the Henry lab (University of Illinois) and are available for download from NCBI SRA database under accession number (SRP114839; Henry et al. 2017). Briefly, the short read transcriptome contains Illumina HiSeq 2500 sequencing results of *Crepidula atrasolea* cDNA library from approximately ∼100-200 embryos from each of the following stages: 1) 2-cell to 25-cell, early cleavage-stage embryos, 2) later cleavage-stage embryos including the formation the 4a-4c micromeres, 3) a mix of more advanced cleavage stages undergoing compaction and initiating gastrulation, 4) early gastrula stage embryos, 5) flattened mid to late gastrula stages undergoing epiboly, 6) embryos undergoing elongation, 7) embryos initiating organogenesis, and 8) more advanced embryos with curved shells.

#### Hybrid transcriptome assembly with both short and long read libraries

A *de novo* transcriptome assembly software capable of combining short and long read data, namely rnaSPAdes (version 3.15.4) (Bushmanova et al. 2019), was used for constructing the hybrid transcriptome. The short read transcriptomes from Henry lab were provided as paired-end libraries, while the Iso-seq transcriptomes were provided as full length libraries with the --pacbio flag. Since the short read library was prepared with Illumina’s TruSeq Stranded mRNAseq Sample Prep kit (Illumina), an additional --ss rf option was specified in the command to specify the strandness of the short read paired-end libraries (first read in pair corresponds to reverse gene strand). In summary, the main command for assembling the hybrid transcriptome is as follows: rnaspades.py −1 ${short_R1} −2 ${short_R2} --ss rf --pacbio ${long_ccs} -o ${result_dir}, where “short_R1” and “short_R2” are paths to the read1 and read2 fastq files from the short read library, “long_ccs” is the path to the Iso-seq CCS reads in fastq format, and “result_dir” specifies the output directory. All commands were executed on the Expanse cluster at the San Diego Supercomputer Center (SDSC). rnaSPAdes returns three transcriptomes with soft-default- and hard-filters, each more stringent in keeping high quality, long length transcripts. We used the default filtered results for subsequent steps as recommended in the rnaSPAdes manual. Overall, there are 848,512 transcripts in this hybrid transcriptome, with a median length of 631 nt.

#### Assessment of completeness of the transcriptome

To assess the quality of the transcriptome, we used BUSCO (version 5.3.2) (Seppey et al. 2019) to score its completeness against Metazoa as well as Mollusca databases (metazoa_odb10 and mollusca_odb10). BUSCO analysis was performed on the SDSC Expanse cluster. The hybrid transcriptome scored high completeness against both databases, with 99.3% for Metazoa and 91.8% for Mollusca. This is markedly increased from the BUSCO completeness score of either the short or long transcriptome alone: the published transcriptome assembly with only the short read library (accession number GFWJ00000000) scored 95.8% against Metazoa and 82.5% against Mollusca; while the Iso-seq clustered transcriptome scored 90.2% against Metazoa and 73.5% against Mollusca.

#### Combining highly similar transcripts

There are highly similar transcripts in the assembled transcriptome that likely represent the same isoform from the same gene, but with small differences due to polymorphism among the non-inbred population used as sample, or technical artifacts during PCR and sequencing. To remove highly redundant transcripts, we ran CD-HIT (version 4.8.1) (Fu et al. 2012) to cluster transcripts in the transcriptome with similarity >95%; only one transcript from each cluster is then kept in the transcriptome. This analysis was done on the SDSC Expanse cluster. The option “-r 0” was used with the cd-hit-est command to perform only +/+ strand alignment during clustering. CD-HIT clustering reduced the total number of transcripts in the transcriptome from 848,512 to 699,466, while maintaining a high BUSCO completeness (99.2% against Metazoan database and 91.7% against Mollusca).

#### Removing foreign transcripts

We used alien index code to remove any foreign transcripts from our sequencing data (Ryan 2014). This was aimed to remove any potential contamination introduced in the RNA extraction and RNA sequencing steps. We used the supplied metazoan protein sequence database (http://ryanlab.whitney.ufl.edu/downloads/alien_index/) as the “non-alien” sequence database. We also used the supplied non-metazoan protein sequence database with the addition of *Diacronema lutheri* as the “alien” sequence database (Nelson et al. 2021). Each transcript is given an alien index score based on the difference between the E-values of the blast to the metazoan and non metazoan databases. We removed transcripts with an alien index of 45 or higher, meaning they were likely to be contaminant non-metazoan sequences.

#### Inferring coding sequences

We inferred the coding sequences (CDSs) in the assembled and cleaned transcriptome using TransDecoder (version 5.5.0) (Haas and Papanicolaou 2015) on SDSC Expanse cluster. “-m 50” option was used with the “TransDecoder.LongOrfs” command to retain only the open reading frames (Orfs) whose protein products are at least 50 amino acids long. The “TransDecoder.Predict” command was then run to predict likely CDSs and their protein/peptide products. CDSs and their products were then filtered to keep only the ones on the sense strand. This resulted in 241,276 sense CDSs and their corresponding proteins/peptides from 195,575 transcripts in the transcriptome. We performed BUSCO analysis on these 195,575 transcripts hosting predicted sense CDSs, and confirmed the persisting high completeness (99.2% against Metazoa and 91.5% against Mollusca databases), while the remaining transcripts in the transcriptome (without long, sense CDS) scored low completeness (6.5% Metazoa and 3.3% Mollusca). The collection of the resulting protein/peptide sequences are used as the proteome for the subsequent analyses.

#### Protein annotations

We used two strategies to annotate proteins and peptides in the proteome constructed per the previous section. First, we scanned the protein/peptide sequences for protein families and domains documented in the InterPro database (Paysan-Lafosse et al. 2023). This was accomplished by running the InterproScan software (version 5.52) (Jones et al. 2014) on the Bridges-2 supercomputing platform at the Pittsburgh Supercomputing Center (PSC). The list of annotation sources used in InterproScan was specified in the command with the -appl option, which include CDD (v3.18), Coils (v2.2.1), Gene3D (4.3.0), Hamap (v2020_05), MobiDBLite (v2.0), PANTHER (v15.0), Pfam (v33.1), PIRSF (v3.10), PIRSR (v2021_02), PRINTS (v42.0), SFLD (v4), SMART (v7.1), SUPERFAMILY (v1.75), and TIGRFAM (v15.0). This approach annotated 126,224 out of the 241,276 entries in the proteome.

As a complementary approach, we blasted each protein/peptide in the *C. atrasolea* proteome against the NCBI Invertebrate Reference Sequence database (RefSeq) (Pruitt et al. 2012). First, a local BLAST database was made for the Invert-RefSeq database on the Bridges-2 cluster (RefSeq database downloaded: May 13, 2022). Next, the *C. atrasolea* proteome was split into three files: two files each containing 100,000 sequences, and the third file containing 41,276 sequences. Pairwise similarity searches were performed for the proteome against the RefSeq invertebrate database using the “BLASTP” command with the following parameters: “-max_target_seqs 10 -outfmt 6 -evalue 1e-5”. Three tabulated output files were generated which were concatenated into a single file. This approach annotated 86,287 of 241,276 entries in the proteome.

### Orthology inference

#### Species proteomes and data curation for orthology inference

We obtained 72 metazoan species proteomes from publicly available databases, which span 13 different metazoan phyla (including 52 molluscan species) and represent multiple sub-phylum-level metazoan clades. We also analyzed 22 biomineral proteomes: 9 of these biomineral proteomes were for species whose proteomes were already included in the 72 proteomes, while the remaining 11 biomineral proteomes are from species without global proteomes, but were kept as standalone single species for orthogroup inference. Sequence accessions for each transcriptome and biomineral proteome were renamed sequentially and include the first letter of their genus name followed by the first few letters of their species names (e.g. Catra_1, Catra_2, Catra_3). Lookup tables were made to match the renamed accession to their original transcriptome accession.

#### Orthofinder2 analysis to identify orthogroups

Orthology inference to identify conserved orthogroups was performed on the previously mentioned input species proteomes using Orthofinder2 (Emms and Kelly 2019), First, an all-vs-all DIAMOND (version 0.9.15) (Buchfink et al. 2015) provided pairwise similarity scores for all sequences, and was used by Orthofinder2 to normalize for gene-length-bias and perform clustering of sequences into orthogroups using the MCL clustering algorithm (van Dongen 2000). Second, sequences found within orthogroups were aligned using MAFTT (version 7.221) (Katoh et al. 2002), and gene tree inference was performed using FastTree (Price et al., 2009) to generate gene trees from multiple sequence alignments. Third, a phylogenetic search database that takes a query sequence and places it into a maximum likelihood gene tree was created using SHOOT (Emms and Kelly 2019).

#### Kinfin analysis to identify lineage-restricted orthogroups

Orthogroups produced by Orthofinder2 were analyzed by Kinfin (version 1.0.3) (Laetsch and Blaxter 2017) to identify metazoan and molluscan orthogroups, as well as lineage-restricted orthogroups at the genus, family, order, class, and phylum levels. First, we use the term “metazoan orthogroup” to refer to orthogroups that have molluscan *and* non-molluscan taxa represented. Second, we use the term “molluscan orthogroup” to refer to orthogroups that only have molluscan taxa found within them. Third, we use the term “lineage-restricted” to refer to orthogroups that contain proteins from more than one species that form a respective monophyletic clade; an exception being for a species-restricted gene which are genes that were not found in any orthogroup. An example of a lineage-restricted gene could be a “molluscan-restricted” orthogroup that contains proteins from more than one class of molluscan taxa or a “gastropod-restricted” orthogroup containing only gastropod taxa. Next, we identified orthogroups for each of the 185 *Crepidula fornicata* SMPs and determined whether their orthogroups were lineage-restricted to phylum Mollusca, class Gastropoda, order Littorinimorpha, family Calyptraeidae, or genus *Crepidula*. Specifically, we first determined which known *C. fornicata* SMPs did not have homologs in other species (classifying them as species-restricted or *fornicata*-restricted SMPs) and we then performed set comparisons to find the difference of SMP orthogroups in higher taxonomic clades.

### Hybridization chain reaction probe design

Following the hybridization chain reaction (HCR) probe design protocol described in Kuehn et al. (2021), we used a custom software (ÖzpolatLab-HCR, 2021) to generate 15-30 DNA oligo probe pairs to *C. atrasolea* messenger RNA (mRNA) for each gene of interest. Using the generated DNA probe sequences, we ordered custom DNA oligos pools (50 pmol DNA oPools Oligo Pool) from Integrated DNA Technologies (Coralville, Iowa, USA). Probes were stored in nuclease-free H_2_O at a concentration of 1 pmol at −80°C.

### Hybridization chain reaction

Embryos were aliquoted into 1.5 ml tubes with their respective probe set and amplifier. Fixed embryos were stored in methanol and were re-hydrated into 5X SSCT buffer following 5 minute 75% methanol 25% 5X SSCT, 50% methanol 50% 5X SSCT, 25% methanol 75% 5X SSCT, and two 100% 5X SSCT washes. Samples were pre-hybridized in 100 μl of probe hyb buffer (30% formamide; # JT4028-1, 5X sodium chloride sodium citrate, 9 mM citric acid (pH 6.0): # C1909, 0.1% Tween 20, 50 μg/ml heparin: Sigma-Aldrich # H3393, 1X Denhardt’s solution: Thermo Scientific # AAJ63135AE, 10% dextran sulfate: Sigma-Aldrich # S4030) at 37°C for 30 minutes. After removing the probe hyb buffer, samples were incubated in 100 ul of the probe solution (1 pmol probe in probe hyb buffer) for 20-24 hours at 37°C. The probe solution was then removed from each sample and washed out with four 5 minute and two 30 minute rinses with probe wash buffer (30% formamide, 5X sodium chloride sodium citrate, 9 mM citric acid (pH 6.0), 0.1% Tween 20, and 50 μg/ml heparin) at 37°C. After washing out the probe, samples were rinsed twice with 5X SSCT for 5 minutes at 20°C. Between these washes, hairpins (H1 and H2) corresponding to the amplifiers used for each tube were snapped separately at 95°C for 90 seconds and then cooled for 30-32 minutes at room temperature before used for the hairpin solution (6 pmol of hairpin solution for each hairpin in 100 ul of amplification buffer per tube). After removing the 5X SSCT solution from each tube, the samples were incubated in the hairpin solution for 22-24 hours at 20°C. Hairpins were then removed by three consecutive 5 minute 5X SSCT washes.

### Negative controls for HCR

We confirmed signal specificity with control samples that were hybridized without a probe set. In the B3 546 fluorophore hairpin-only control veliger embryos, we noted very faint labeling in the putative larval kidney, velum, and ocellus. We also noted faint background labeling of the putative larval kidney and ocellus in our veliger B1 647 controls.

### Hoechst staining

Immediately after the HCR (detailed above), samples were stained for either 2-3 hours at ∼20°C or overnight at 4°C with 0.001 mg/ml hoechst in 5X SSCT (5X sodium chloride sodium citrate, and 0.1% Tween 20 in nuclease-free H_2_O). Samples were then rinsed with 5X SSCT three times for 5 minutes at 20°C. Samples were stored at 4°C for up to a week.

### Mounting and imaging

Following Hoechst and/or WGA staining, individual specimens were mounted in either 80% Glycerol (Promega H5433) 20% 5X SSCT or 2% methyl cellulose in 5X SSCT (for posterior imaging) on Rain-X-coated (ITW Global Brands, Houston, TX) 22 x 50 mm glass coverslips for fluorescent imaging on a Zeiss LSM 700 on the 20x objective with an AxioCam HRm camera. Fluorescent images were acquired in ZenBlack (Zeiss), then processed in ImageJ and Adobe Photoshop (Adobe Inc., California, USA). Samples for dark field images were mounted as described above, and imaged on a Zeiss Axio Imager M2 on the 10x objective with an AxioCam 506 Color camera. Darkfield images were acquired in ZenBlue (Zeiss), then processed in Helicon Focus (HeliconSoft (RRID: SCR_014462) and ImageJ. Figures were created in Adobe Illustrator (Adobe Inc., California, USA).

## Results

### Complementary models of *Crepidula* for studying molluscan biomineralization

We identified two *Crepidula* species, *C. fornicata* and *C. atrasolea,* that together serve as complementary models to study shell biomineralization (Fig. 1A). In a previous study focusing on *Crepidula fornicata* (Batzel et al. 2022), we identified over 185 SMPs in the adult shell and 20 SMPs differentially expressed in the adult mantle tissue. Of these 20 differentially expressed adult SMPs, at least 10 showed shell field expression during larval development in traditional chromogenic *in-situ* hybridization. This study demonstrated that some adult SMPs are also involved in larval shell production. *C. fornicata* was ideal for that initial study because its large size facilitates protein extraction from shells and mRNA extraction from dissected mantle tissue. However, *C. fornicata* is not ideal for studying shell development, since embryos are not available year round. This is due to its delayed metamorphosis (at least 2-3 weeks; (Pechenik 1984), long generation time ( >1 year), and seasonal reproduction (Lyons and Henry, 2022). To avoid these drawbacks, we have begun building resources and tools for *C. atrasolea* as an alternative and complementary research organism for studying shell development (Henry et al. 2017; Lyons and Henry 2022). *C. atrasolea* is a direct-developer with a generation time of ∼4 months, and with a modest colony can provide embryos year-round (Henry et al. 2017; Lyons and Henry 2022). While *C. fornicata* is an excellent model for transcriptomic and proteomic approaches, *C. atrasolea* is the ideal model for studying shell development. In this study we use our *C. fornicata* SMP dataset to examine the conservation of SMPs across different clades, and *C. atrasolea* to further examine the spatial-temporal expression of conserved SMPs during larval development.

Despite their life history differences, we found that the two species are very similar from spiral cleavage through organogenesis (Fig. 1B). Shell field similarities (e.g. morphological development and expansion of tissue from the posterior) are observed at the ovoid and organogenesis stages ( Fig. 1B). Morphological differences arise later in larval development; the planktonic feeding larvae of *C. fornicata* contain larger velar lobes, to the pre-hatch veliger of *C. atrasolea* (Fig. 1B). Major shell morphological differences, including color, shape, and size, are most visible in adults (Fig. 1C).

### Characterization and timing of shell development in *Crepidula atrasolea*

To establish *C. atrasolea* as a model research organism for shell development in gastropods, we generated a detailed staging scheme examining stage-specific morphological features of shell development (Fig. 2). Refining the published staging scheme in (Henry et al. 2017), we based our stages on development at 20°C with a focus on morphological changes to the shell field, the embryonic and larval shell-forming tissue. We found that wheat germ agglutinin (WGA), a cell membrane marker with a high affinity to N-acetylglucosamine and N-acetylneuraminic acid residues (Mishkind et al. 1982; Fodero et al. 2001; Baddeley et al. 2011), clearly marks the shell field throughout larval shell development.

**Figure 2:**
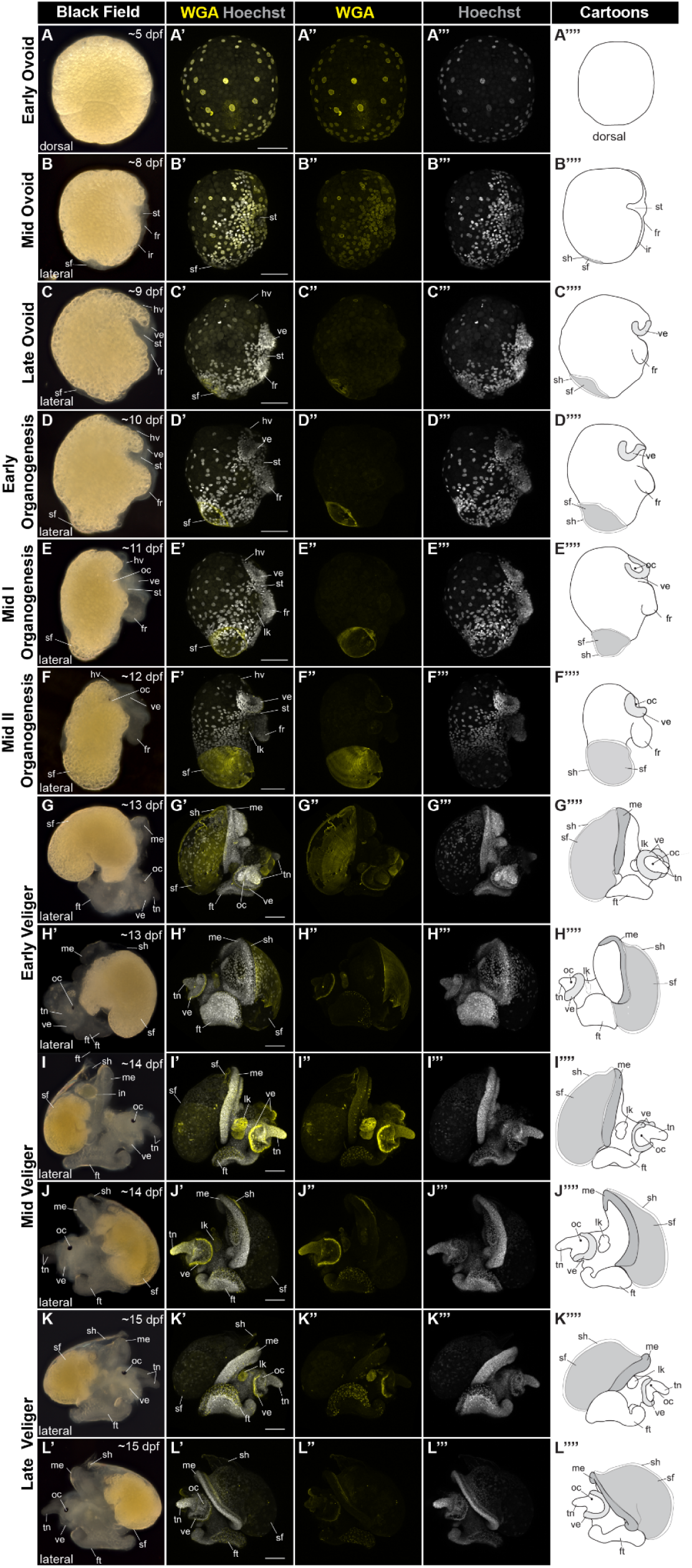
Characterization and timing of shell development in *Crepidula atrasolea*. Dark field and confocal images of fixed *C. atrasolea* embryos from approximately 8 days post fertilization (dpf) to approximately 15 dpf at 20°C. Fixed embryos were stained with wheat germ agglutinin (WGA) and imaged at the 20x objective on a confocal microscope. Cartoons highlight changes in shell morphology during development. In early and mid ovoid staged embryos (A-B), WGA stains nuclear membranes throughout the embryo. WGA begins to mark the shell field starting at the late ovoid stage (C). WGA continues to mark the developing shell throughout the organogenesis stages (D-F). WGA also marks the velar lobes, larval kidney, and foot in organogenesis and veliger staged embryos (F-L). Hoechst is shown in gray and WGA in yellow. fr, foot rudiment; ft, foot; hv, head vesicle; lk, larval kidney; in, intestine; ir, intestinal rudiment; oc, ocellus; sf, shell field; st, stomodeum; tn, tentacles; ve, velar lobes. Scale bars each represent 100 μm.

Organic shell secretions–as assayed by WGA staining–in the shell field are first detected during ovoid stages (8-9 dpf). Larval structures, including the foot rudiment, stomodeum, shell field, and intestinal rudiment, were first visible in mid ovoid embryos at ∼8 dpf, when elongation along the anterior-posterior axis is apparent (Fig. 2A). During the mid ovoid stage (∼8 dpf), the posteriorly-located shell field is translucent and rounded. Translucent ventral structures, including the velar lobes and foot rudiment, become more prominent. At the mid ovoid stage, WGA intensely stains the outlines of cell nuclei throughout the embryo and very weakly stains the developing shell field in the posterior terminus (Fig. 2A’). The late ovoid stage (∼9 dpf) is distinguished by the presence of two ciliated velar lobes, a lengthened foot rudiment, and a flattened shell field along the dorsal-posterior end of the embryo (Fig. 2B). At this stage, glycoprotein secretion from the mantle cells into the extracellular matrix is first visible, as indicated by bright WGA staining (Fig. 2B’).

During organogenesis stages (10-12 dpf), larval shell accretion corresponds to elongation along the anterior-posterior axis (Fig. 2C: Fig. 2E). In the early organogenesis stage (∼10 dpf), the shell field begins to diverge from the midline for the first time, as it rounds out in the posterior of the embryo (Fig. 2C). WGA staining is visible in the larval shell and putative larval kidney during early organogenesis (Fig. 2C’). At the mid I organogenesis stage (∼11 dpf), the shell field elongates along the anterior-posterior axis, larval organs continue to develop, and pigmented ocelli are visible (Fig. 2D). The larval shell field thickens and expands in the mid II organogenesis stage (12 dpf), and the pigmented ocelli are pronounced (Fig. 2E). Increased WGA staining highlights shell field expansion during mid I organogenesis (Fig. 2D’) and mid II organogenesis (Fig. 2E’).

In veliger stages (∼13 dpf to ∼20 dpf), the larval shell is fully formed, and larval organs are well developed (Fig. 2F: Fig. 2K). At the early veliger stage, an enlarged foot extending ventrally along the body, enhanced velar lobes, and paired tentacles are also present (Fig. 2F and Fig. 2G). The WGA staining of glycoproteins in the shell field diminished during the veliger stages. At mid veliger stages (∼14 dpf) the shell is angled upwards (Fig. 2H and Fig. 2I), and then flattens out to cover the entire animal during late veliger stages (Fig. 2J and Fig. 2K). The yolk reserves begin to diminish during the veliger stages until they are no longer visible in the hatched juvenile at ∼21 dpf.

In summary, we characterized the detailed morphological changes of *C. atrasolea* during larval shell development with high temporal resolution, and identified WGA as a marker for the shell field and nuclei membrane.

### Nearly all SMPs in the *C. fornicata* shell proteome were found in orthogroups

Molluscan biomineral proteomes have been shown to contain a small repertoire of highly conserved SMPs and a larger overall fraction of novel SMPs, also referred to as species-restricted SMPs (Clark 2020). In order to determine the evolutionary origins of SMPs and their placement into orthogroups, we performed orthology inferences on proteins from *C. fornicata* and *C. atrasolea* proteomes, as well as proteomes of 85 additional species spanning the metazoan phylogeny. Orthology inference was performed using Orthofinder (Emms and Kelly 2019); the 185 SMPs previously identified in the shell proteome of *C. fornicata* (Batzel et al. 2022) were included in the analysis as part of the *C. fornicata* proteome. Briefly, Orthofinder defines the pairwise similarities between all sequences in the provided proteomes based on length-corrected alignment scores, then uses the resulting similarities to cluster the sequences into orthogroups; genes contained in each orthogroup are considered orthologs (Emms and Kelly 2019).

Using these methodologies, we asked whether *C. fornicata* SMPs have orthologs in other species, or whether most are novel, species-restricted SMPs. Orthorfinder assigned 89.5% (n = 2,860,915 of 3,197,660) of the input proteins to 165,643 orthogroups. We examined the orthofinder results with a focus on *C. fornicata* SMPs and determined that 95.6% (n = 177 of 185) of the SMPs were assigned to 150 orthogroups (Fig. 3). We found that of the 95.6% of SMPs found in orthogroups, that 43.2% (n = 80) of these SMPs were in orthogroups that contain taxa from at least one molluscan species and one non-molluscan species, which we classify as a “metazoan orthogroup”. Additionally, we determined that the remaining 52.4% (n=80) of SMPs contain taxa from at least one molluscan species, which we classify as a “molluscan orthogroup”. On the other hand, our results showed that only 4.4% (8 of 185) of SMPs were not found in any orthogroup, suggesting that few species-restricted SMPs exist in the shell proteome. Taken together, these results demonstrate that the majority of SMPs have conserved orthologs while few SMPs are actually novel to *C. fornicata*. This contradicts the generally-held notion that an animal’s biomineral proteome is dominated by mostly novel genes.

**Figure 3:**
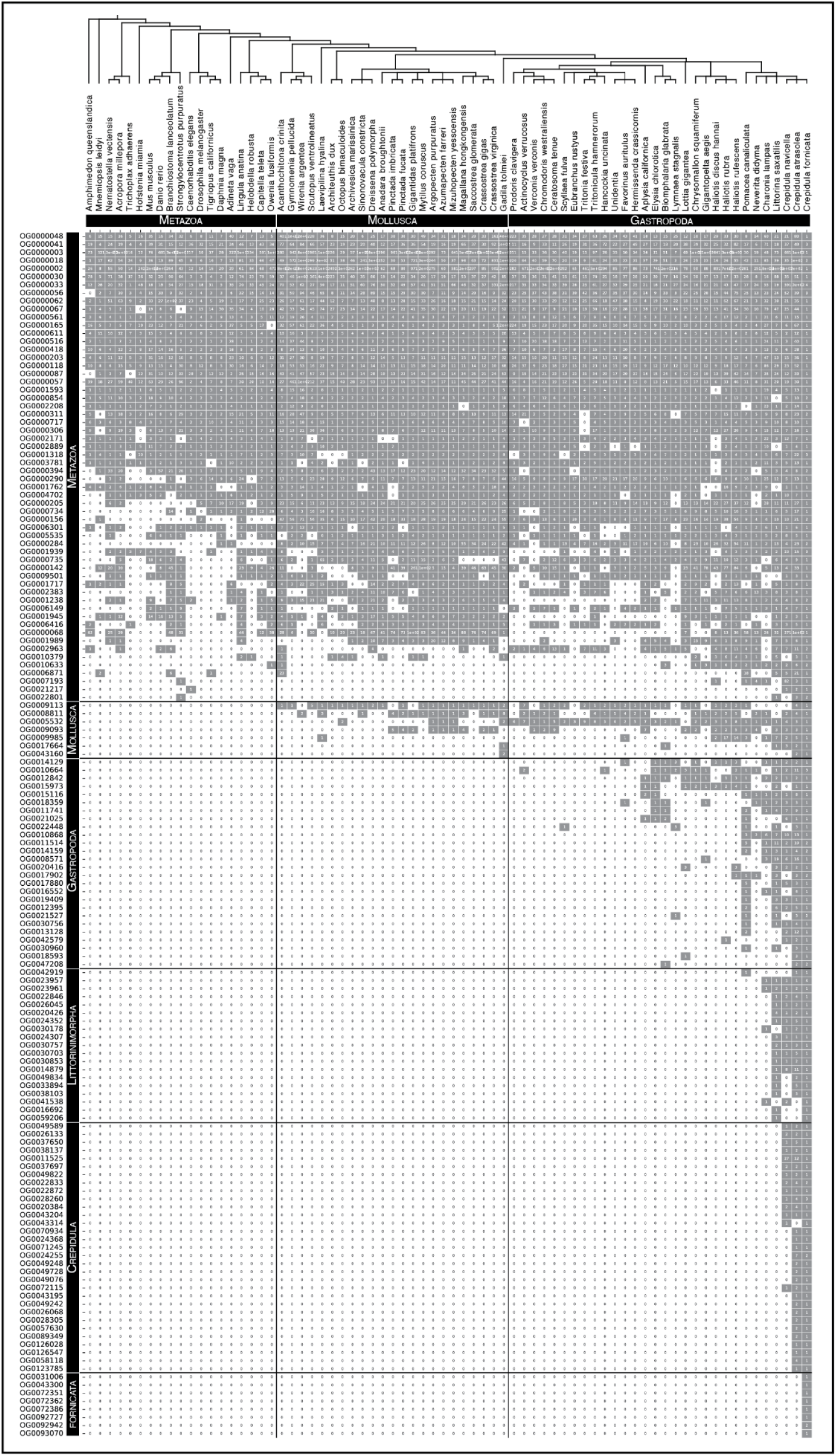
Shell matrix proteins of *Crepidula fornicata* clustered into orthogroups. Rows represent SMP orthogroups that were generated through ortholog inference techniques, and ordered based on the taxonomic level they are found to be lineage-restricted. Columns depict counts of proteins per taxon with their order (tree depicted above columns) based on current understandings of their phylogenetic positions (Kocot et al. 2011; Laumer et al. 2015; Laumer et al. 2018; Cunha and Giribet 2019). Gray boxes indicate presence of at least one protein from a taxon in an orthogroup. White boxes indicate that no protein was found for a respective taxon in an orthogroup.

### Lineage-restricted orthogroups account for at least half of all shell proteome SMPs

In order to understand how shell morphological characteristics arise within species, we need to identify their evolutionary origins and the taxonomic level at which an SMP is lineage-restricted. We define lineage-restricted orthogroups as orthogroups that have more than one taxon represented within their respective clade-level groupings, unless the orthogroup contains only one taxon, in which case we refer to it as a species-restricted gene. An example of the former being a molluscan-restricted orthogroup which contains species from at least two different classes (e.g. a bivalve and gastropod ortholog present in an orthogroup). We focused our analyses on molluscan orthogroups, which were found to represent 52.4% of all SMPs. These molluscan orthogroups were further sub-divided into multiple lineage-restricted SMP groups: 4% (n = 7) Molluscan-restricted; 18% (n = 33) Gastropod-restricted; 12% (n = 22) Littorinimorpha-restricted; 19% (n = 33) *Crepidula*-restricted (Fig. 4). We observed that *Crepidula*-restricted SMP orthogroups were responsible for nearly one fifth of all lineage-restricted SMP orthogroups, and that of the two other *Crepidula* species in the dataset, *C. atrasolea* had orthologs for 92.9% (n = 172 of 185) of all SMPs, compared to 73.4% (n = 136 of 185) for *C. navicella.* These results demonstrate that SMPs can be classified into lineage-restricted groups, and that closely related species share a similar repertoire of SMPs. The ability to classify SMPs at different taxonomic levels will allow us to generate experimental hypotheses regarding the functional role that lineage-restricted SMPs have on the evolution of the molluscan shell.

**Figure 4.**
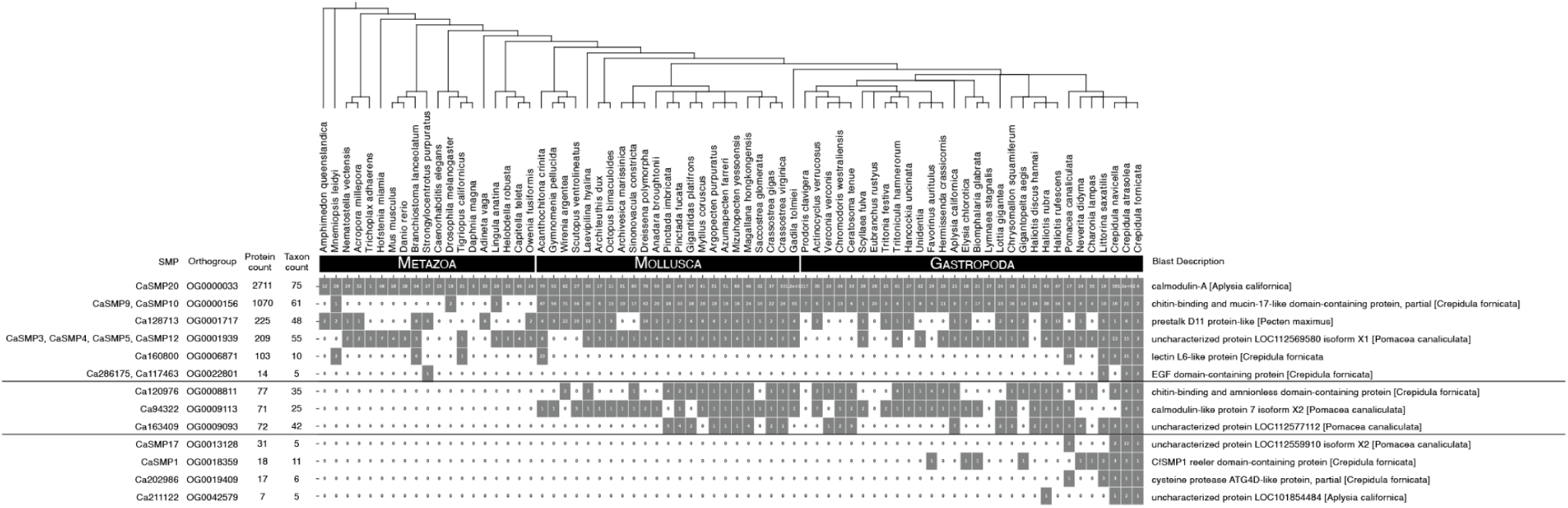
Lineage-restricted shell matrix proteins selected for HCR in *Crepidula atrasolea*. Rows represent SMP orthogroups that were examined by HCR in *C. atrasolea.* They are ordered based on the taxonomic level they are found to be lineage-restricted (*Metazoa*-restricted, *Gastropoda*-restricted, and *Crepidula*-restricted). Columns depict counts of proteins per taxon with their order (tree depicted above columns) based on current understandings of their phylogenetic positions (Kocot et al. 2011; Laumer et al. 2015; Laumer et al. 2018; Cunha and Giribet 2019). Gray and white boxes indicate presence or absence of respective species within SMP orthogroups.

### Lineage-restricted orthogroups selected for Hybridization Chain Reaction (HCR)

Distinct expression domains of SMPs in the mantle have been observed across the Mollusca (Jolly et al. 2004; Marie et al. 2012; Herlitze et al. 2018), and were hypothesized to contribute to shell microstructure and morphological variation between different clades (Jackson 2021). We asked whether genes in different lineage-restricted orthogroups exhibit different expression domains in *C. atrasolea* shell field, and whether putatively annotated uncharacterized proteins show expression in the shell field. For HCR in *C. atrasolea*, SMPs were selected at different lineage-restricted levels, prioritizing SMPs that were previously found differentially expressed in the mantle compared to the head, foot, and gill tissues. In total, 18 SMPs in 13 different orthogroups were examined by HCR in *C. atrasolea* (Fig. 4). These orthogroups contain 11 metazoan-restricted, 3 molluscan-restricted, and 4 gastropod-restricted SMPs. To annotate orthogroups, BLAST searches for *C. fornicata* SMP homologs were performed against GenBank and their best reciprocal BLAST hit descriptions were used as putative annotations of the orthogroup As a result, 4 orthogroups returned BLAST hit descriptions of uncharacterized proteins, while the remaining 9 orthogroups returned descriptions of proteins that have been previously implicated in biomineralization including chitin-binding, calmodulin, lectin-like, and cysteine-protease to name a few.

### Shared shell field-specific expression of SMP1 in *C. atrasolea* and *C. fornicata* embryos

To confirm that gastropod-restricted SMPs have comparable expression patterns in *C. fornicata* and *C. atrasolea*, we compared larval expression of shell matrix protein 1 (SMP1), the most upregulated SMP in the adult *C. fornicata* mantle, in the two species. We found exclusive shell field expression of SMP1 in both *C. fornicata* and *C. atrasolea* (Fig. 5). The CaSMP1 homolog we found contains an extracellular matrix-binding reeler domain, which is also found in *C. fornicata*. Based on the morphology of the velar lobes and foot rudiment, we identified late ovoid and late organogenesis stage embryos as the most comparable stages between the two species (Fig. 1C). Using HCR, we found that expression of CfSMP1 and CaSMP1 was confined to cells within the shell field at the ovoid stages (Fig. 5A and Fig. 5C), and a stronger ring of expression was seen in the mantle edge during organogenesis (Fig. 5B and Fig. 5D). We used WGA to stain the larval shell field in *C. fornicata and C. atrasolea* embryos (Fig. 5B: Fig. 5D). WGA staining of the shell field in *C. fornicata* was very faint at the ovoid stage (Fig. 5A’’’) and most prominent at the organogenesis stage (Fig. 5B’’’). Meanwhile, WGA staining was bright in both ovoid (Fig. 5C’’’) and organogenesis staged embryos of *C. atrasolea* (Fig. 5D’’’). In both species, the WGA staining encompasses the SMP1 expressing cells (Fig. 5A: Fig. 5D). Together, SMP1 and WGA showed a more dorsal-lateral positioned, and flatter, shell field in *C. fornicata* embryos compared to the larval shell field of *C. atrasolea*.

**Figure 5.**
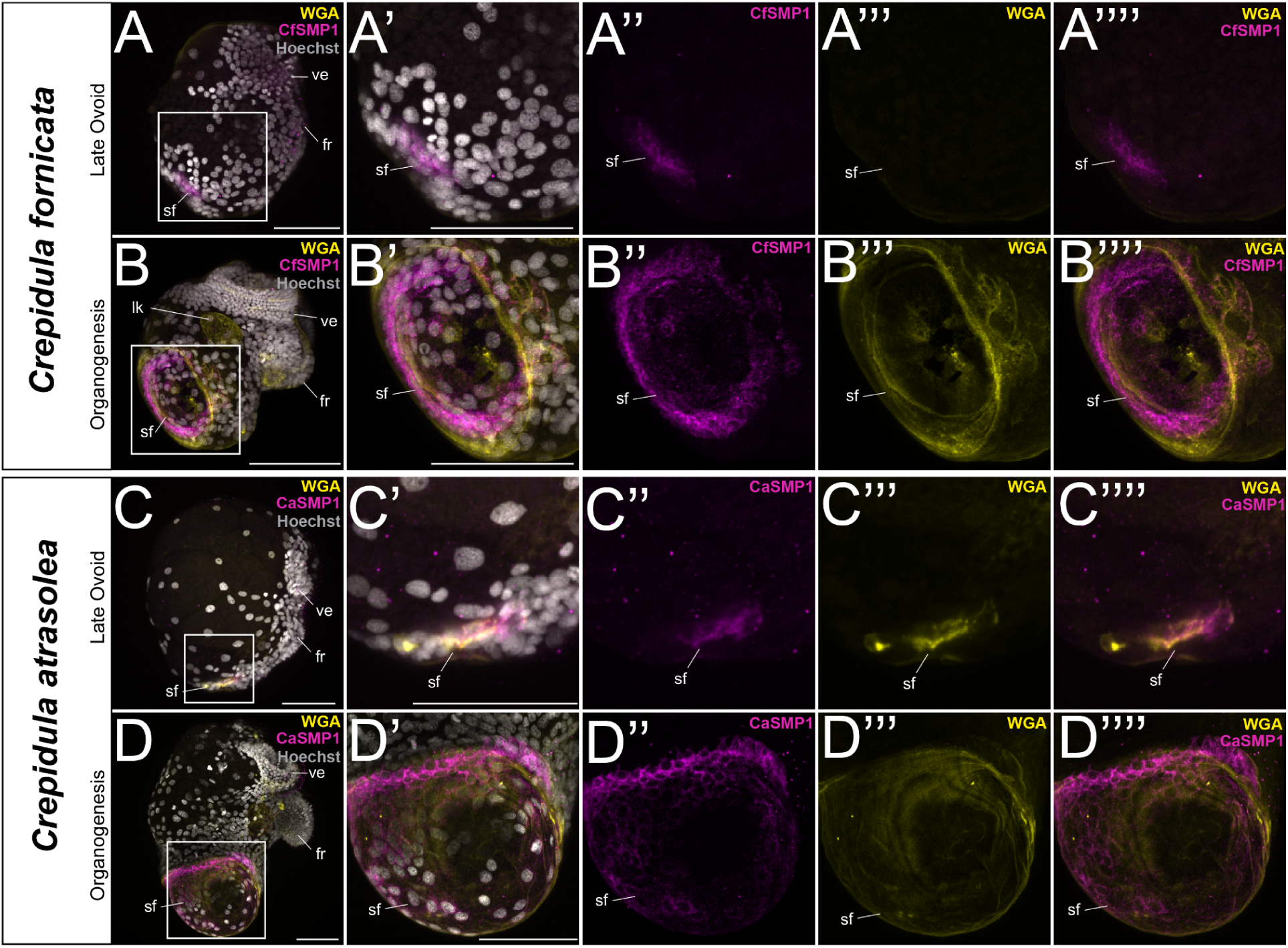
Shell field specific expression of shell matrix protein 1 mRNA in *Crepidula fornicata* and *Crepidula atrasolea* embryos. Using hybridization chain reaction (HCR), mRNA expression was detected for SMP1 in both species (A-D). Wheat germ agglutinin (WGA) was used to mark the larval shell. CfSMP1 is expressed throughout the shell field in late ovoid staged *C. fornicata* embryos (A), and more intensely around the mantle edge during organogenesis (B). Similar expression patterns are seen for CaSMP1 in late ovoid staged (C) and organogenesis staged *C. atrasolea* embryos (D). Hoechst is shown in gray, SMP1 in magenta, and WGA in yellow. fr, foot rudiment; ft, sf, shell field; ve, velar lobes. Scale bars represent 100 μm.

In both colorimetric *in situ* hybridization and HCR, CaSMP1 was detected in the mantle edge. However, colorimetric *in situ* hybridization failed to capture CaSMP1 expression throughout the shell field. In *C. fornicata*, we found that the HCR pattern revealed a finer cellular resolution of CfSMP1 expression in the mantle edge compared to that of colorimetric in-situs. Given these limitations to colorimetric detection, we decided that HCR was the preferred method for examining the expression profiles of *C. atrasolea* SMPs. It also has the added advantage of probe multiplexing, allowing us to examine the expression patterns of multiple genes at once.

### 12/18 SMPs examined are expressed in larval *C. atrasolea* mantle tissue

To identify SMPs expressed in the shell field of *C. atrasolea*, we characterized the spatial-temporal expression of 18 different SMPs at the late organogenesis and veliger stages. Out of the 18 genes we examined, 12 were expressed in the shell field, and five (CaSMP1, CaSMP10, CaSMP9, CaSMP17, and Ca211122) were exclusively expressed in the shell field (Fig. 6 and Fig. 7). Many of the SMPs we examined were expressed in other tissues outside of the shell field, including in the putative statocyst and the velar lobes. We found two general SMP expression patterns in the shell field: SMPs with broader expression throughout the shell field (Fig. 6), and SMPs with more specific expression in the mantle edge (Fig. 7).

**Figure 6.**
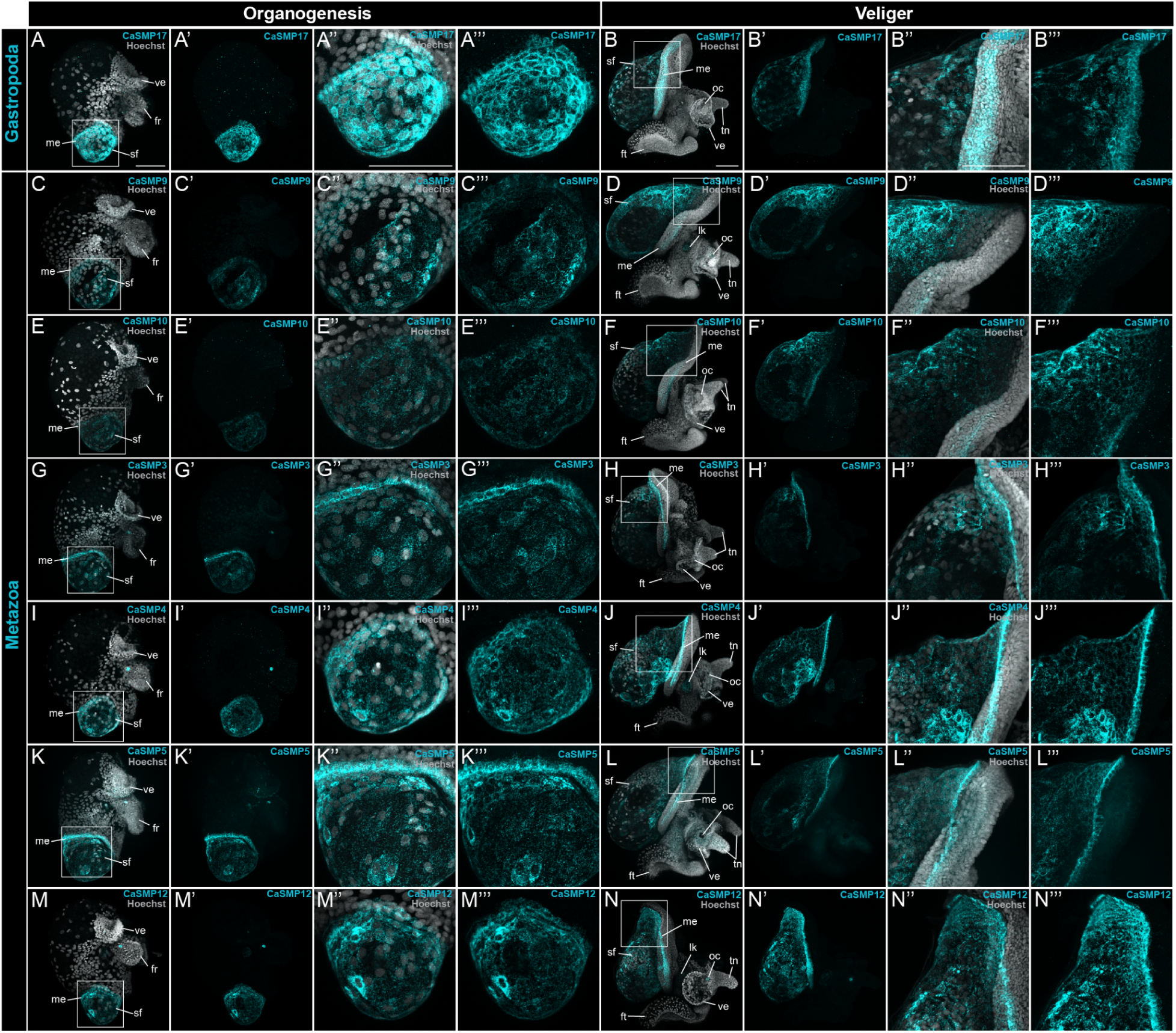
Broad shell field SMP expression in *Crepidula atrasolea* at organogenesis and veliger stages. Using hybridization chain reaction, mRNA expression was detected for CaSMP17 (A and B), CaSMP9 (C and D), CaSMP10 (E and F), CaSMP3 (G and H), CaSMP4 (I and J), CaSMP5 (K and L), and CaSMP12 (M and N). Hoechst is shown in gray and each SMP in cyan. ft, foot; sf, shell field; tn, tentacles; ve, velar lobes. Scale bars in each represent 100 μm.

**Figure 7.**
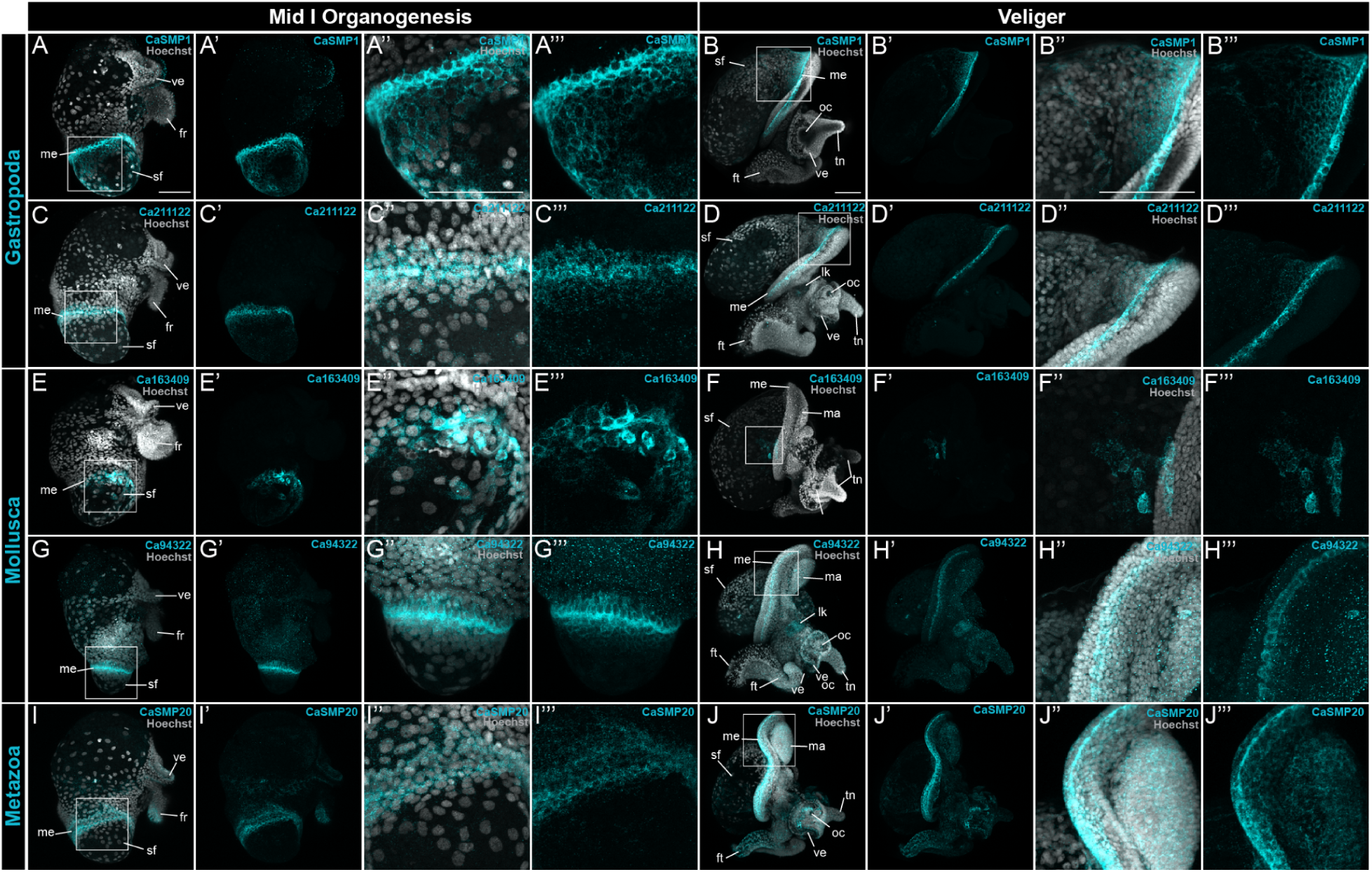
Restricted mantle edge SMP expression in *Crepidula atrasolea* at organogenesis and veliger stages. Hybridization chain reaction detected mRNA expression of CaSMP1 (A and B), Ca211122 (C and D), Ca163409 (E and F), Ca94322 (G and H), and CaSMP20 (I and J) in organogenesis and veliger staged embryos. Hoechst is shown in gray and each SMP in cyan. ft, foot; sf, shell field; tn, tentacles; ve, velar lobes; lk, larval kidney. Scale bars each represent 100 μm.

### 7/12 SMPs are broadly expressed in the shell field

#### Uncharacterized gastropod-specific CaSMP17

We identified CaSMP17 as an uncharacterized gastropod-specific SMP (Fig. 3). CaSMP17 expression was confined to columnar and cuboidal cells in the shell field during organogenesis (Fig. 6A) and veliger stages (Fig. 6B). In organogenesis-staged embryos, expression was detected in every cell of the shell field (Fig. 6A’’). In veliger-staged embryos, CaSMP17 expression shifted to a patchy distribution throughout shell field cells, and in cells of the mantle edge (Fig. 6B’’’).

#### Metazoan mucin-like

Mucin and mucin-like genes are cell surface glycoproteins (Strous and Dekker 1992) with a cytoplasmic C-terminus and an extracellular, glycosylated, N-terminus, which are separated by a transmembrane region (Carraway and Hull 1991). Mucins are secreted by epithelial cells and aid biomineralization of vertebrate bones, teeth, and cartilage (Midura and Hascall 1996). Mucin-like genes have been previously identified in molluscan mantle tissue, and are hypothesized to aid shell biomineralization (Marin et al. 2000; Marin and Luquet 2005; Zeng and Guo 2022). In *C. atrasolea*, we identified two mucin-like genes, CaSMP9 (Fig. 6C and D) and CaSMP10 (Fig. 6E and Fig. 6F), that each contain two chitin binding protein domains. Both of these genes are in the metazoan orthogroup (Fig. 3). We found that these genes are expressed throughout shell field cells during organogenesis (Fig. 6C and Fig. 6E) and veliger stages (Fig. 6D and Fig. 6F). While expression is distributed throughout the mantle tissue, the brightened signal is detected in the dorsal-anterior cells of the veliger shell field (Fig. 6D’’’ and Fig. 6F’’’).

#### Uncharacterized metazoan SMPs

We found four uncharacterized genes, CaSMP3, CaSMP4, CaSMP5, and CaSMP12 in the metazoan orthogroup (Fig. 3). We found that these uncharacterized SMPs had patchy expression throughout the shell field and in a subset of cells along the mantle edge at both the late organogenesis and veliger stages (Fig. 6G: Fig. 6N). In the late organogenesis stages, expressing cells were also located in the putative statocyst (as seen in Fig. 6G’ and Fig. 6I’). While there was some individual variability in the intensity of signal, we also observed bright expression in a band of squamous cells in the mantle edge during organogenesis (Fig. 6G’’ and Fig. 6K’’). In the veliger stages, patchy expression persisted throughout the shell field, with a brighter, continuous band of cells labeled in the mantle edge (Fig. 6H; Fig. 6J; Fig. 6L; Fig.6N). CaSMP4 positive cells were highly concentrated in an anterior-ventral patch within the shell field where the shell is curved to one side (Fig. 6J’).

### 5/12 SMPs have restricted mantle edge expression

#### Reeler domain-containing CaSMP1

We identified CaSMP1 as the *C. atrasolea* homolog of CfSMP1, and as a gastropod lineage restricted SMP (Fig. 3). In mid I organogenesis stage embryos, we observed expression throughout the shell field with a brighter band of expression along the mantle edge (Fig. 7A). In veliger stage embryos, expression becomes more restricted to the mantle edge (Fig. 7B), with diminishing expression in shell cells further from the mantle edge (Fig. 7B’’).

#### Uncharacterized gastropod-specific Ca211122

We identified Ca211122 as a gastropod-specific uncharacterized protein with restricted shell field expression in *C. atrasolea* (Fig. 3; Fig. 7C; Fig. 7D). In late organogenesis stage embryos, Ca21122 expression was confined to a continuous band of cells along the mantle edge, with speckled expression throughout the shell field (Fig. 7C). During the veliger stages, Ca211122 persisted in the mantle edge and was weakly expressed in the dorsal-anterior portion of the shell field (Fig. 7D).

#### Uncharacterized molluscan-specific Ca163409

We identified Ca163409 as an uncharacterized molluscan-specific SMP (Fig 3) with a narrow range of expression along the mantle edge (Fig. 7E and Fig. 7F). In mid organogenesis stage embryos, Ca163409 expression was visible in a subset of ventral-laterally positioned shell field cells (Fig. 7E’’). The most prominent expression was detected adjacent to the mantle edge (Fig. 7E’’’). In veliger stage embryos, Ca163409 was expressed in a small discrete group of cells located within the anterior portion of the shell field, adjacent to the mantle edge (Fig. 7F’’). We noted variability in the number of shell field cells labeled between specimens, and also noted anterior expression in the head of one individual.

#### Calmodulin family SMPs

Calmodulin and calmodulin-like are highly conserved eukaryotic calcium-binding proteins with EF-hand domains. They belong to a superfamily involved in numerous signaling pathways including cellular calcium metabolism (Means and Dedman 1980; Perochon et al. 2011; O’Day et al. 2015), cell proliferation (Agell et al. 2002), and epithelial cell differentiation (Rogers et al. 2001). The structure and expression of calmodulin-like has been well characterized in the adult mantle epithelium tissue of the pearl oyster *Pinctada fucata* (Li et al. 2004; Li et al. 2005; Li et al. 2006; Fang et al. 2008). In *P. fucata*, calmodulin-like and calmodulin proteins have different subcellular localizations within the mantle epithelium; calmodulin-like proteins are concentrated only in the cytoplasm while calmodulin is found in both the nuclei and cytoplasm (Fang et al. 2008). While calmodulin and calmodulin-like are hypothesized to play different roles in molluscan biomineralization, the function of these proteins remains uncertain. In gastropods, multiple copies of calmodulin superfamily genes have been identified in the tropical abalone *Haliotis asinina (Jackson et al. 2006; Jackson et al. 2007)*, the pacific abalone *Haliotis discus* (Lim et al. 2018), the California sea hare *Aplysia californica* (Swanson et al. 1990), and the periwinkle, genus *Littorina (Simpson et al. 2005)*. In *H. asinina* larval and juvenile stages, calmodulin expression is localized to ectodermal cells along the mantle edge (Jackson et al. 2006; Jackson et al. 2007), suggesting calmodulin’s involvement in gastropod shell growth via calcium secretion and regulation.

We identified two calmodulin family genes, Ca94322 and CaSMP20, with two different expression patterns within the mantle edge during organogenesis and veliger stages (Fig. 7G: Fig. 7J). Ca94322 is a calmodulin-like 7 protein in the molluscan orthogroup (Fig. 3). Ca94322 expression was strongest in one ring of cells around the mantle edge of the larval shell field (G’’’). Ca94322 expression diminished in cells further from the mantle edge (Fig. 7G’’’). At the veliger stage, the most intense expression was seen within a continuous band of Ca94322-positive cells between the outer and inner mantle folds (Fig. 7H and Fig. 7H’’). During the veliger stages, Ca94322 expression expanded to the anterior portion of the foot, and diminished levels were detected throughout the head of the embryo (Fig. 7H).

We also found a calmodulin A, CaSMP20, in the metazoan orthogroup (Fig. 3). During organogenesis, CaSMP20 expression was most prominent in two rings of cells along the mantle edge of the larval shell field (Fig. 7I’’). Expression was also detected in the posterior terminus of the foot rudiment, and faintly throughout the velar lobes and anterior portion of the body (Fig. 7I). During the veliger stages, CaSMP20 expression expands throughout the anterior-ventral portion of the animal, with pronounced expression in the mantle edge (Fig. 7J and Fig. 7J’’). Faint foci of expression were observed in the posterior portion of the shell field (Fig. 7J’). The mantle edge-specific patterns of *C. atrasolea* calmodulin family genes were similar to those reported in *P. fucata (Li et al. 2004)* and *H. asinina* (Jackson et al. 2006; Jackson et al. 2007).

### Co-expression of CaSMP3 and CaSMP20 mRNA reveals three distinct mantle cell populations during larval development

To determine if different regions of SMP expression (e.g. broader vs. mantle edge specific domains) correspond to distinct mantle cell populations, we examined the spatial-temporal expression of CaSMP3 and CaSMP20 throughout larval shell development (Fig. 8). We confirmed that CaSMP20 and CaSMP3 have two discrete domains and one overlapping region of expression in late ovoid to veliger staged embryos (Fig. 8). We identified three shell field cell populations during larval development: a CaSMP3+ population in the shell field, a CaSMP20+ population lining the edge of the shell field, and a CaSMP3+/CaSMP20+ population in between. Specifically, at the late ovoid stage, posteriorly located CaSMP3 positive cells were surrounded by a fainter ring of CaSMP20 expression, with a small population of cells expressed both CaSMP3 and CaSMP20 (Fig. 8A: Fig. 8C). At the early organogenesis stage three cell populations are discernible by a patch of CaSMP3 expression in posterior end, a ring of CaSMP3 and CaSMP20 co-expression around the shell field, and a second, outer ring of CaSMP20 expression in the mantle edge (Fig. 8D: Fig. 8F). At this stage, these three cell populations were most visible on the posterior end (Fig. 8F’ and Fig. 8F’’). In mid organogenesis embryos, CaSMP3 expressing cells were located throughout the shell field, CaSMP3 and CaSMP20 co-expressing cells were concentrated to a band along the mantle edge, and CaSMP20 expressing cells were situated along a second band above the CaSMP3 and CaSMP20 positive cell population (Fig. 8G’ and Fig. 8H’). At the veliger stage, we saw a laterally asymmetric distribution of shell field cell populations (Fig. 8I and Fig. 8J). On the left lateral side there were two prominent bands of CaSMP20 expression within the mantle edge (Fig. 8J’’’). CaSMP3 and CaSMP20 were co-expressed in cells of the second, posterior-most band (Fig. 8J’’). The third cell population, marked by CaSMP3 expression, was located throughout the shell field (Fig. 8J’’’’). In contrast, there was only one, thicker band of CaSMP20 positive cells on the right lateral side (Fig. 8I’’’), with the CaSMP3 and CaSMP20 co-expressing cell population restricted to a smaller patch of the mantle edge (Fig. 8I’’).

**Figure 8.**
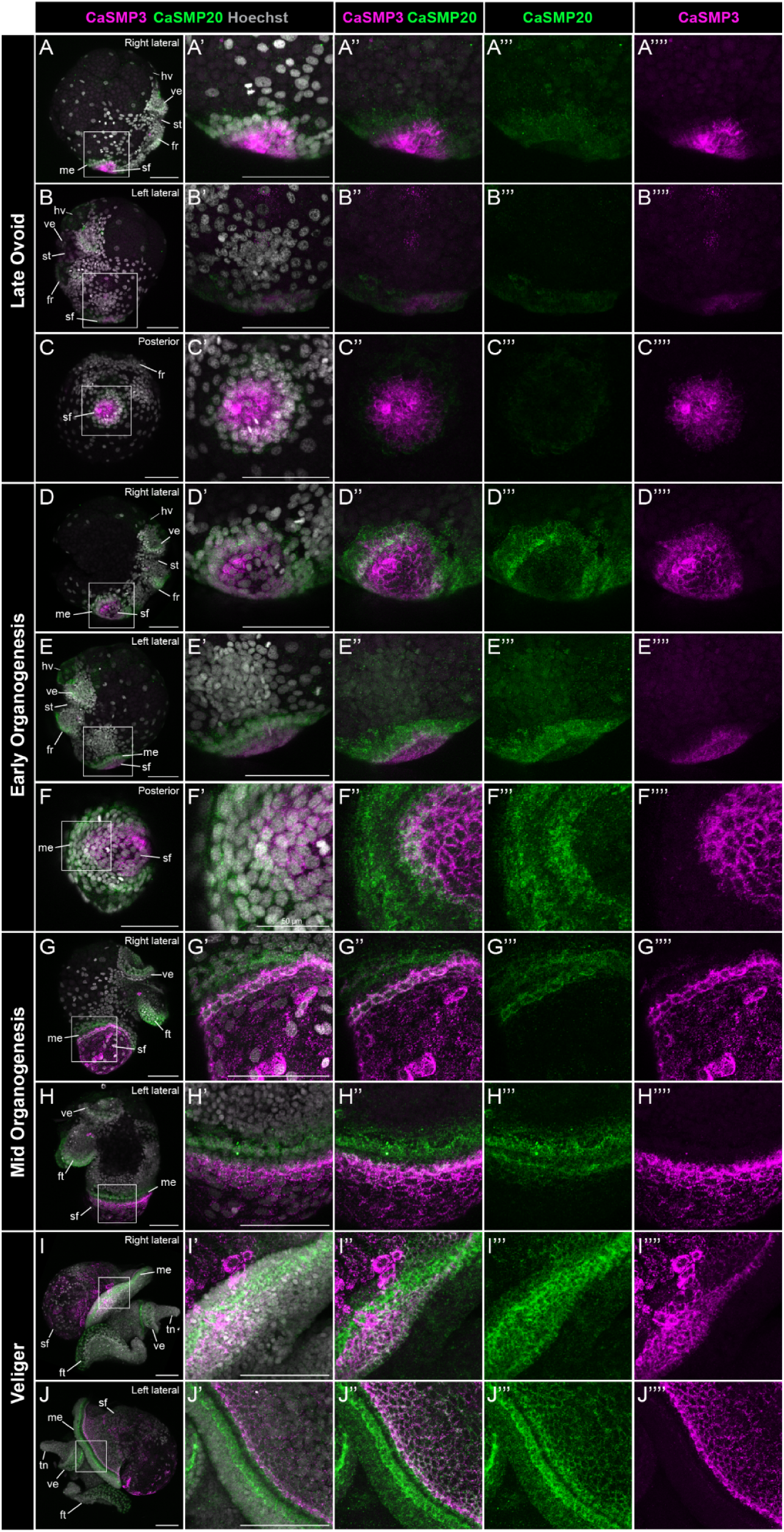
Co-expression of CaSMP3 and CaSMP20 mRNA throughout embryonic shell development in *Crepidula atrasolea*. Right lateral (A), left lateral (B), and posterior views (C) of late ovoid embryos show co-expression of CaSMP3 (magenta) and CaSMP20 (green) in the shell field. These two expression domains continue during early organogenesis (D-F) and mid organogenesis (G-H). In the veliger stages, laterally asymmetric expression is most apparent in the shell field and mantel edge (I-J). Hoechst is shown in gray, CaSMP3 in magenta, and CaSMP20 in green. ft, foot; sf, shell field; tn, tentacles; ve, velar lobes. Scale bars each represent 100 μm unless otherwise specified.

We also observed larval expression of CaSMP3 and CaSMP20 outside of the shell field, with higher levels of CaSMP20 expression in anterior and ventral tissues compared to CaSMP3 (Fig. 8). From the late ovoid to veliger stages, the CaSMP20 signal was detected faintly in patches throughout the embryo and more prominently in three non-shell field structures: the posterior edge of the foot rudiment, the head vesicle, and the velar lobes. (Fig. 8A: Fig. 8F). Apart from the shell field, CaSMP3 was only expressed in the larval statocysts (Fig. 8A: Fig. 8H) Taken together, CaSMP3 and CaSMP20 exhibit specificity to the shell field and mark novel SMP expressing cell populations that have yet to be examined in other molluscs.

### Summary of regionalized SMP shell field expression

Through our spatial-temporal expression analysis of 18 SMPs, we identified 12 SMPs from a range of lineage-specificities that are expressed in the shell field (Fig. 9A). We initially detected two domains within the shell field: the broader shell field, and the mantle edge (detailed above). Within these general domains, we found 5 distinct expression patterns (Fig. 9B), and 3 distinct cell populations that are regionalized within the shell field (Fig. 9C). Based on the spatial-temporal location of these cell populations, we identified three primary regions of the shell field: the outer mantle edge, the inner mantle edge, and the broader shell field (Fig. 9C).

**Figure 9.**
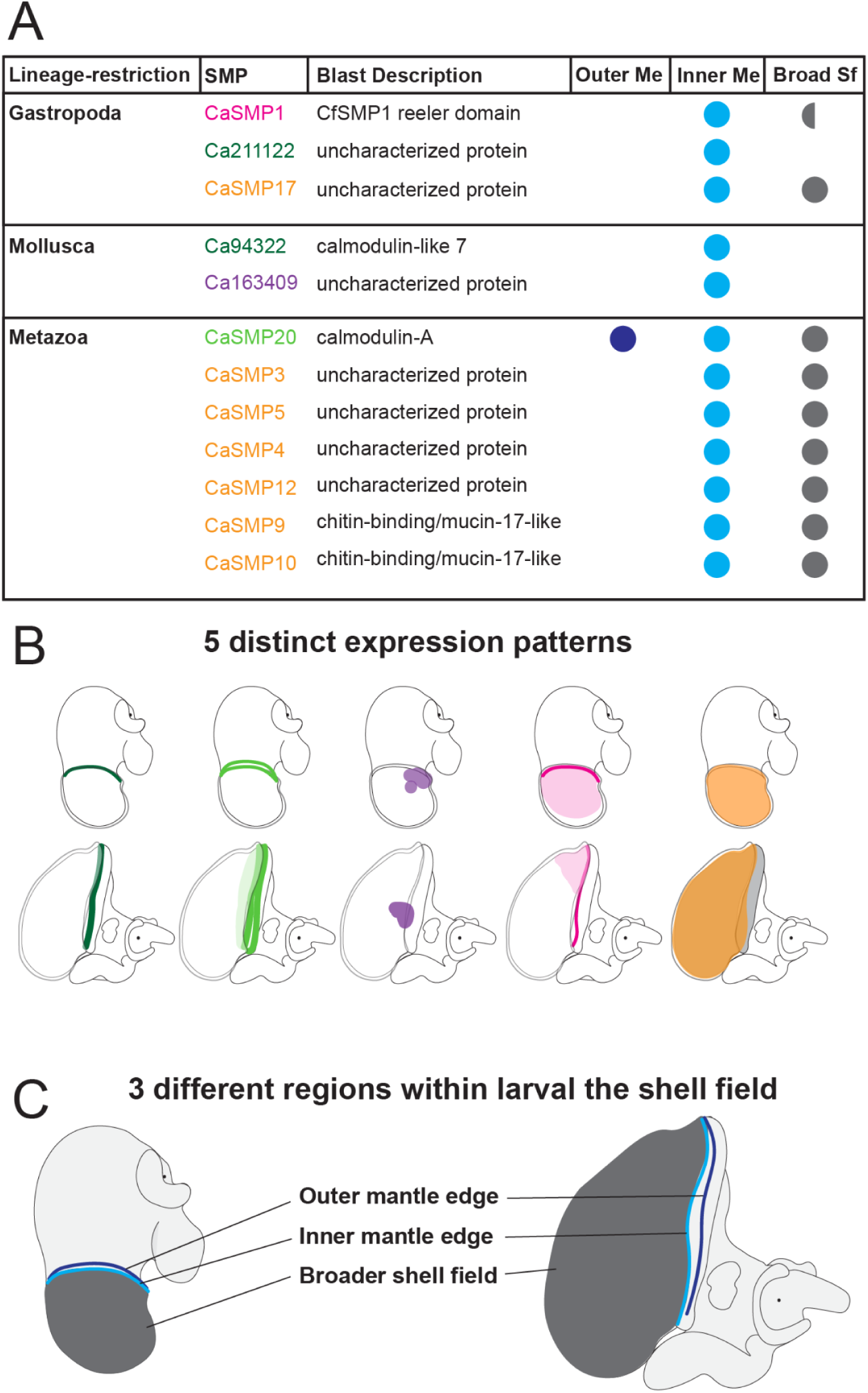
Summary of regionalized SMP expression during larval shell development in *Crepidula atrasolea*. Summary table showing the lineage-restriction of each SMP, the best blast hit description, and shell field region(s) of expression (A). Circles in (A) indicate presence or absence of expression in the outer mantle edge (outer Me), inner mantle edge (inner Me), and/or the broader shell field (broad Sf). Half circles indicate presence of expression in one stage (late organogenesis or veliger) but not both. Cartoons depicting 5 different SMP expression patterns within the shell field of organogenesis and veliger stage *C. atrasolea* embryos (B). The SMPs are color coded in (A) based on their expression patterns shown in (B). Cartoons showing regionalization of the shell field into 3 potential zones of biomineralization in late organogenesis and veliger stage *C. atrasolea* embryo (C). These zones are highlighted in different colors: outer mantle edge in dark blue, inner mantle edge in light blue, and broader shell field in dark gray (C).

## Discussion

### *Crepidula atrasolea* and *Crepidula fornicata* as models for molluscan biomineralization

We showed that transcriptomic and proteomic work in *C. fornicata* can be directly integrated into our understanding of shell biomineralization in *C. atrasolea*. From our orthofinder analysis of *C. fornicata* SMPs, we discovered many SMP orthologs with differing evolutionary ages that are conserved in *C. atrasolea*. Additionally, we found comparable expression patterns of SMP1 in the shell field of both species. Given its specificity to the gastropod lineage, SMP1 is likely a conserved gastropod shell field marker. By directly comparing SMP1 expression with WGA staining, we were able to visualize glycoprotein secretion from mantle epithelial cells into the extracellular matrix in both species. The larval shell field of other gastropods could be examined using WGA staining in combination with HCR for SMP1 homologs.

Based on the lineage-specificity of the 185 adult *C. fornicata* SMPs, we selected 18 SMPs for spatial-temporal characterization in *C. atrasolea*. From this analysis in *C. atrasolea*, we described shell field modularity and identified 3 shell field cell populations. We also found 12 SMPs that are expressed in the adult stage of *C. fornicata* (Batzel et al., 2022), that are also expressed during larval shell development in *C. atrasolea*, suggesting shared SMP expression in adult and larval mantle epithelia. Leveraging data from both species, we provide a comprehensive analysis of larval SMPs and build a foundation of knowledge in the genetically tractable molluscan model, *C. atrasolea*. Future efforts in *C. atrasolea* can use our findings to determine the SMP cell type specificity and function in molluscan shell biomineralization.

### Adult SMPs are expressed during larval shell development in *C. atrasolea*

We found 12 adult SMPs from *C. fornicata* that are expressed during larval shell development in *C. atrasolea*, suggesting shared SMP expression in adult and larval mantle epithelia. This observation opposes previous transcriptome and shell proteome comparisons that found drastically different SMPs expressed in larval vs. adult bivalve shells (Zhao et al. 2018; Carini et al. 2019; Cavallo et al. 2022). Zhao et al. (2018) identified only 4 SMPs that were shared between larval and adult bivalves (*Pinctada fucata* and *Crassostrea gigas*), and Cavallo et al. (2022) identified only 1 out of 5 adult SMPs with larval expression in the Antarctic clam *Laternula elliptica*. Meanwhile, we found that 12/18 of the adult SMPs we examined via HCR were expressed at larval stages. Our results may highlight differences in bivalve vs gastropod biomineralization, and/or suggest that spatial-temporal expression data is required to further validate transcriptomic and proteomic findings.

Since the candidate SMPs we selected for HCR were both upregulated in adult *C. fornicata* mantle tissue (Batzel et al. 2022) and conserved in *C. atrasolea*, the SMPs with larval expression likely contribute to core biomineralization processes that are required throughout mantle maturation. It is possible that SMPs without shell field expression serve a function unique to adult biomineralization. To determine the complete repertoire of SMPs shared during larval and adult biomineralization, future work should screen all 185 adult SMPs for larval expression patterns in both *C. fornicata* and *C. atrasolea*.

### *Crepidula* SMPs have regionalized shell field expression

Molluscan mantle modularity has been widely observed in adult tissue (Sudo et al. 1997; Jolly et al. 2004; Takeuchi and Endo 2006; Zhang and Zhang 2006), but few studies have focused on larval tissue (Hohagen and Jackson 2013). During larval shell development of *Lymnaea stagnalis*, Herlitze et al. (2018) detected regionalized expression of SMPs to specific zones of the shell field. Similarly, in *C. atrasolea* we found distinct regions of SMP expression: broad expression throughout the shell field, and more restricted expression in the mantle edge. Through HCR and confocal imaging, we were able to attain a high cellular and subcellular resolution of the mRNA localization, and identified additional domains of SMP expression within the mantle tissue. We observed that HCR was much more sensitive than traditional colorimetric *in situ* hybridization, especially in non-mantle edge regions, and we recommend that this method be used in other species.

Different cell populations of the shell field have been identified in the bivalve *Crassostrea gigas* (Liu et al. 2020) and in multiple gastropod species (Nederbragt et al. 2002; Hohagen and Jackson 2013; Herlitze et al. 2018; Johnson et al. 2019). For instance, a band of cells with high proliferation rate occupies the outermost portion of the shell field, which we refer to as the “mantle edge”, also known as the “leading edge” or “aperture growth zone” (Johnson et al. 2019). We examined the spatial-temporal expression of multiple SMPs within the same individual to identify distinctive SMP-expressing cell populations. We identified CaSMP3 expressing cells located throughout the shell field, CaSMP3+ and CaSMP20+ cells in the inner mantle edge, and CaSMP20 expressing cells in the outer mantle edge. This study is the first to identify SMP-expressing cell populations, and provides a comprehensive characterization of SMP expression within the mantle edge of the shell field. We also confirmed the conservation of these cell populations in the mantle tissue of *C. fornicata* larvae. To further characterize these cell populations, and determine cell fate specification programs, these SMP markers could be used to identify specific mantle cell populations within single cell seq datasets, for example. Because SMP3 and SMP20 are widely conserved across metazoans, their homologs could be examined in the single-cell seq data sets of other molluscan species.

### Lineage-restricted SMPs partly explain mantle modularity

Novel shell characteristics could have arisen through the expression of either novel SMPs or conserved SMPs in different regions of the mantle. We hypothesized that highly conserved, metazoan SMPs might be involved in fundamental biological processes in biomineralization (Gilbert et al. 2022) and would show broader shell field expression compared to more recently-diverged, lineage-restricted SMPs, which might show restricted expression in the mantle edge where shell growth occurs. We found that the majority of broadly-expressed *C. atrasolea* SMPs in the shell field were from metazoan orthogroups (n= 6/7), and the majority of SMPs with a more restricted mantle edge expression were lineage-restricted to Mollusca or Gastropoda (n= 4/5). This may suggest that lineage-restricted SMPs tend to be expressed in more specific regions of the mantle tissue. There are exceptions to this trend; for example, SMP20 is a metazoan SMP that was found restricted to the mantle edge, and CaSMP17 is a gastropod-restricted SMP, with a broader shell field expression domain. To thoroughly test the relationship of evolutionary age of SMPs to their spatial expression in the shell field, we would need to characterize the expression of all lineage-restricted and metazoan SMPs in *C. atrasolea*. Additionally, in the future we can test these hypotheses through CRISPR-Cas9 mediated knock-outs in *C. fornicata* and *C. atrasolea* (Perry and Henry 2015; Henry et al. 2017; Lyons and Henry 2022), which we have now established as complementary models for understanding molluscan shell biomineralization. We can determine if perturbation of species-restricted SMPs would disrupt biomineralization in *C. atrasolea*, and what consequences that would have for shell morphology and microstructure.

### Previous approaches underestimate evolutionary conservation of SMP orthogroups

Understanding the phylogenetic relationships of SMPs is essential for tracing the evolution of biomineralization in marine invertebrates. Within the past two decades, a surprising outcome from proteomic investigations of biomineral structures is the preponderance of lineage-restricted shell matrix proteins (Clark 2020). Examples in bivalves include 52% in *Pinctada maximus*, 54% in *Mya truncata*, and 66% *Mytilus edulis* of SMPs had no BLAST hit ((Arivalagan et al. 2017). These observations have been informed in part by BLAST pairwise similarity *E*-value scores against large sequence databases, under the assumption that high pairwise similarity-scoring SMPs may have evolved more recently because they lack homologous reciprocal BLAST hits. Indeed, close examination of the sequences of lineage-restricted SMPs have shown they contain long stretches of repetitive low complexity regions (Marie et al. 2013), which are generally a hallmark of recently-diverging genes (Toll-Riera et al. 2012). Likewise, in our previous study we took a BLAST approach and determined that 29% of *C. fornicata* SMPs had no best-reciprocal BLAST hit against sequences in GenBank, and that many of these SMPs contain long stretches of repetitive low complexity domains (Batzel et al. 2022).

However, we argue here that a lack of reciprocal BLAST hit is not always a reliable measure of lineage restriction or recent divergence events, as it can be an artifact due to homology detection failure in heuristic BLAST searches, which result in high numbers of false-negatives, especially in short and rapidly evolving genes (Weisman et al. 2020). This is due to the BLAST algorithm placing greater emphasis on pairwise-similarity scores that are directly correlated with gene-length and without regard to variable sequence evolution. This detection failure suggests that species-restricted SMPs may have homology outside of their own lineages, but due to their highly divergent sequences, their degree of similarity does not meet a certain threshold by BLAST standards. A second explanation for homology detection failure is the underrepresentation of species in GenBank. We observed a large number of *C. fornicata* SMPs are shared by two other *Crepidula* species: *C. navicella* and *C. atrasolea*. These two species were found to have orthologs for 92.9% (*C. atrasolea*) and 73.4% (*C. navicella*) of *C. fornicata* SMPs. Without these two species, the number of *fornicata*-restricted SMPs would increase from 4.8% to 23.7%, the latter being similar to our previous findings that 29% of *C. fornicata* SMPs had no BLAST hits. This discrepancy could be explained by underrepresented taxa, especially from related species, being absent in BLAST databases like GenBank. Previously, *C. atrasolea* and *C. navicella* species transcriptome assemblies were not searchable through GenBank, leaving open the possibility that other underrepresented taxa, in addition to gene-length-bias, may result in homology detection failure. These results underscore the importance of broad taxon sampling, and data accessibility, for future studies of molluscan biomineralization genes.

### Orthology inference of SMP orthogroups is improved using a cluster-based approach

While BLAST approaches are generally used to understand sequence homology between two sequences from different species, they are highly ineffective at extending their similarity score to orthologous genes from multiple species (Emms and Kelly 2019). Recently, clustering-based algorithmic approaches for inferring gene orthology have emerged, which do not rely solely on pairwise similarity scores, but instead apply weighted connections to pairwise similarity scores in order to form orthologous clusters (Li et al. 2003; Emms and Kelly 2015; Emms and Kelly 2019). Moreover, elimination of gene-length bias from BLAST searches has the potential to transform our understanding of SMP diversity by reclassifying previously annotated lineage-restricted SMPs. This is evident in the fact that a typical BLAST search of an SMP containing low-complexity regions against GenBank might find multiple regions of alignment, but those regions produce high pairwise similarity scores that are above a user-determined BLAST cutoff, thus returning no significant best hits. In contrast, a clustering approach like Orthofinder applies normalization to the pairwise similarity score to account for sequence length of the query and length of its hits, ensuring that even *distantly* related sequences receive equivalent scores compared to the best-scoring sequences from *closely* related species (Emms and Kelly 2015). This suggests that rapidly evolving regions in lineage-restricted SMPs, which are susceptible to homology detection failure, may actually have a homolog in a last common ancestor when accounting for gene-length bias. The consequence of this would be that BLAST approaches have underestimated the total number of SMP orthogroups. This was observed in our analysis of *C. fornicata* SMPs, as we recovered 34 more SMP orthologs with the orthofinder approach compared to the previous BLAST approach. Thus we recommend that future studies of molluscan SMPs incorporate a cluster-based approach.

## Acknowledgements

We thank the Hamdoun Lab (UC San Diego) for the use of their confocal, and the Ozpolat Lab (Washington University) and the Katz Lab (UMass Amherst) for advice about adapting HCR to Crepidula embryos. This work was funded by a National Science Foundation Faculty Early Career Development (CAREER) Award [1943606] and a National Institute of General Medical Sciences Maximizing Investigators’ Research Award (MIRA) [R35GM133673] to DCL.

